# Olfactory neuroepithelium biomarker platform: a proof from Alzheimer’s disease

**DOI:** 10.64898/2026.10.18.759991

**Authors:** Koko Ishizuka, Guanqun Shang, Saba Malik, Carolyn Doty, Shrey Dave, Nicholas Rowan, Yukiko Lema, Andrew Lane, Akira Sawa

**Affiliations:** Departments of Psychiatry, Johns Hopkins University School of Medicine, Baltimore, MD, 21287, USA; Departments of Otolaryngology-Head and Neck Surgery, Johns Hopkins University School of Medicine, Baltimore, MD, 21287, USA; Departments of Biomedical Engineering, Neuroscience, Johns Hopkins University School of Medicine, Baltimore, MD, 21287, USA; Departments of Pharmacology, Johns Hopkins University School of Medicine, Baltimore, MD, 21287, USA; Genetic Medicine, Johns Hopkins University School of Medicine, Baltimore, MD, 21287, USA; Department of Mental Health, Johns Hopkins Bloomberg School of Public Health, Baltimore, MD, 21205, USA

## Abstract

Unlike other diseases where target organs for biopsy are accessible, surrogate platforms are warranted for biomarkers of neuropsychiatric disorders due to the inaccessibility of the brain. We demonstrate a non-invasive platform combining a nasal soft-brush swab with single-cell Western blotting for neuron-specific protein analysis. Applied to Alzheimer’s disease targeting phosphorylated tau, this demonstrated superior diagnostic capacity compared with blood-based approaches, indicating that the olfactory neuroepithelium is a sensitive biomarker platform.

## Main

Biospecimens from living patients are essential for investigating disease mechanisms, identifying biomarkers, and developing targeted therapies. However, obtaining brain specimens from living individuals remains a major challenge in brain disorders. Therefore, approaches that enable neuron-specific molecular analysis in living patients are highly desirable^1,2^. While blood-based biomarkers represent a promising trend, they indirectly evaluate brain-derived molecular and cellular materials, frequently affected by peripheral factors^3-6^. In contrast, the olfactory neural epithelium (OE), which contains mature olfactory neurons, has emerged as an accessible surrogate that may capture neuronal signatures more directly^2,7-10^. Alzheimer’s disease (AD), the most common cause of dementia, exemplifies the need for accessible neuronal biomarkers. Positron emission tomography (PET) imaging and cerebrospinal fluid (CSF) analysis of amyloid-β (Aβ) and tau^11-13^ provide high diagnostic accuracy but are limited by cost, accessibility, and invasiveness. Recently, blood-based biomarkers have also become available^14^. In the present study, we developed a non-invasive, needle-free platform that combines nasal brush swab with single-cell Western blotting (scWB) to enable neuron-specific protein and post-translational modification analyses at single-cell resolution in living patients. As a proof of concept, we applied this platform to AD and evaluated its diagnostic performance relative to blood-based biomarkers.

First, we developed a non-invasive method to collect olfactory neuronal samples by using nasal brush swab at the dorsoposterior region of the nasal septum (**Fig. 1A**). This anatomical site contains the OE enriched with mature olfactory neurons, as confirmed by histological and molecular characterization (**Fig. 1B, C**). Immunostaining of the swabbed tissue from this site identified cells spanning the olfactory neuronal lineage, including keratin 5 (KRT5)-positive horizontal basal cells, βIII-tubulin-positive immature olfactory neurons, and olfactory marker protein (OMP)-positive mature olfactory neurons, as well as surrounding keratin 18 (KRT18)-positive sustentacular cells (**Fig. 1B**). Consistent with these findings, qPCR analysis of RNA isolated from nasal brush samples from the same site specifically detected expression of *OMP* (**Fig. 1C**). Immunostaining of nasal brush samples further confirmed the presence of mature olfactory neurons at the protein level (**Fig. 1D**). Together, these results demonstrate that non-invasive nasal brush swabs can successfully collect mature olfactory neurons.

**Figure 1.**
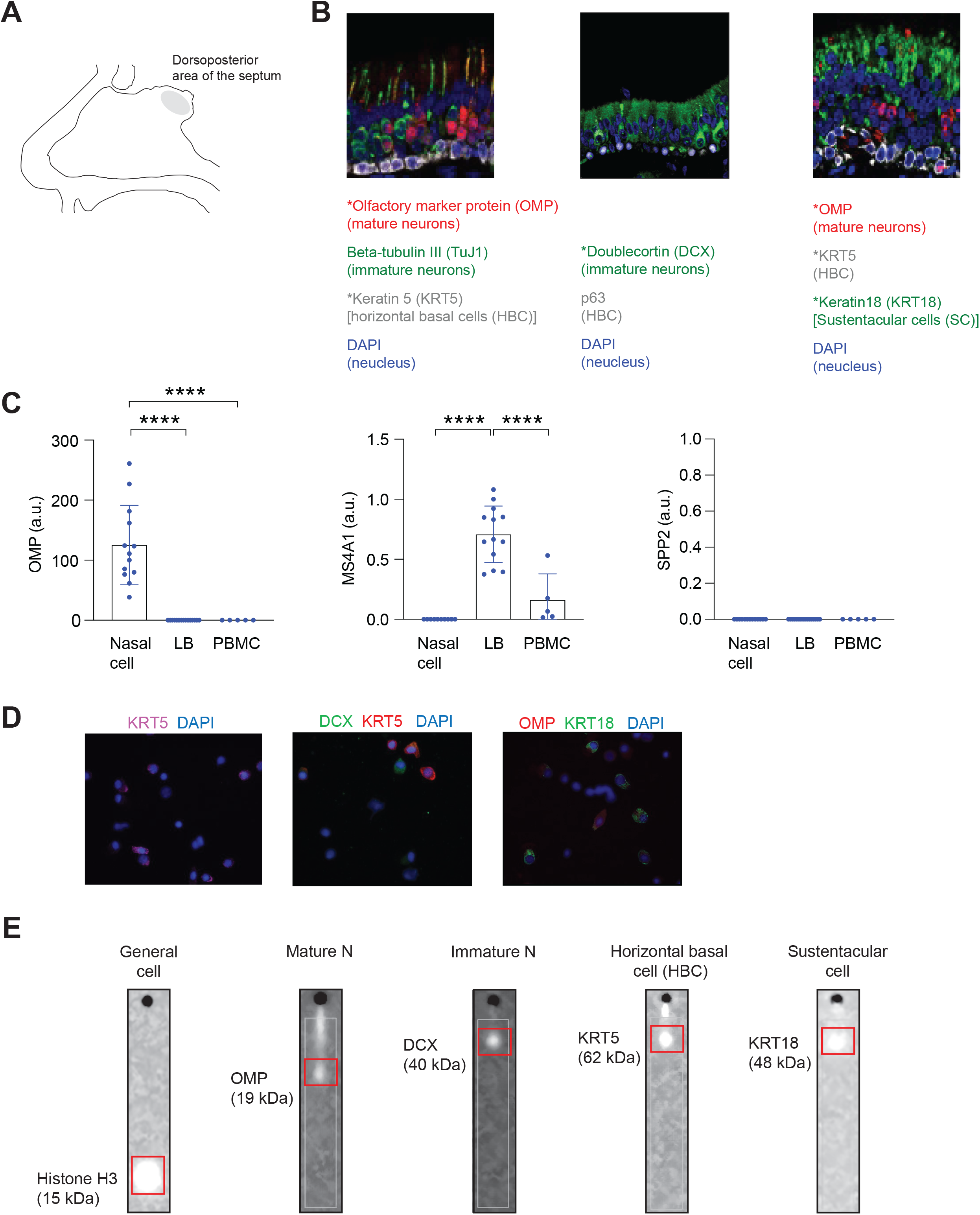
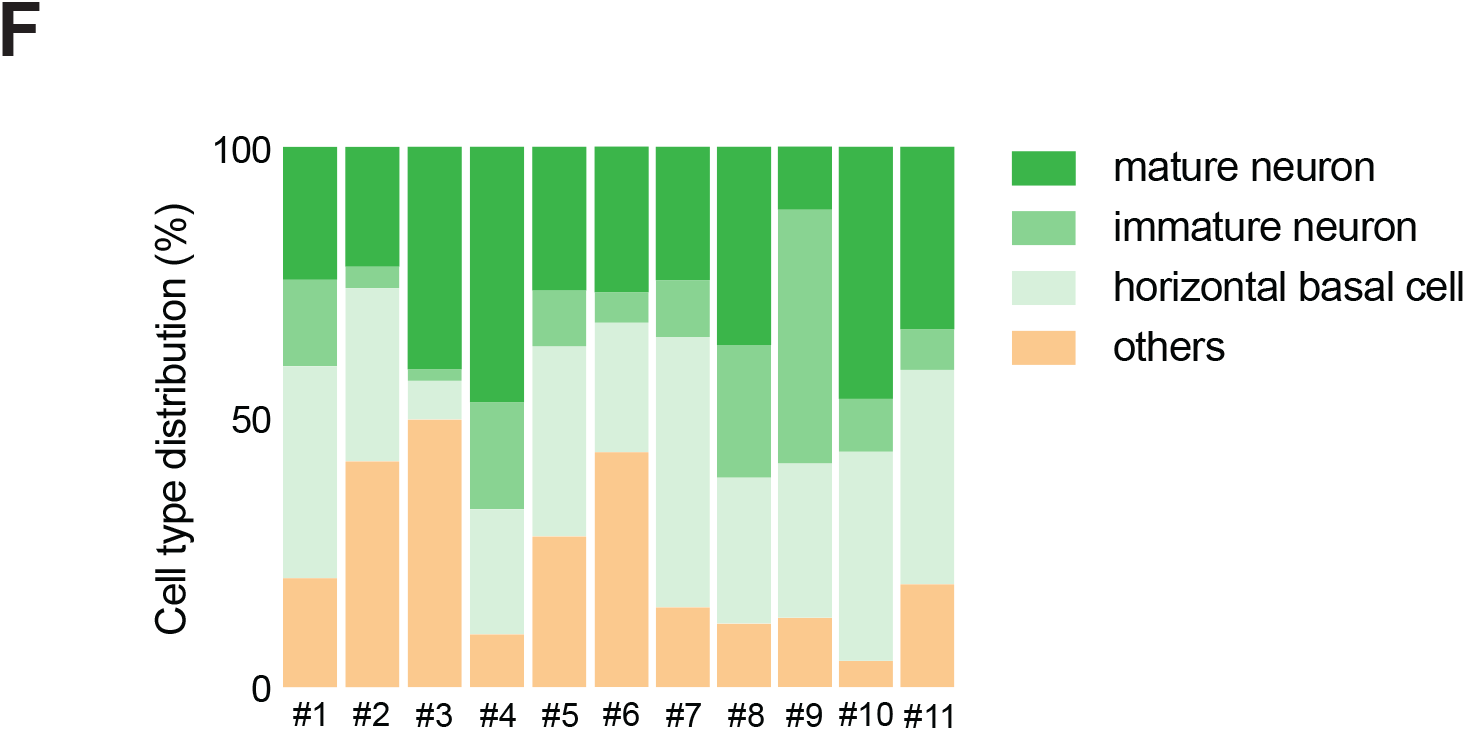
Non-invasive collection of mature olfactory neurons for single-cell level biomarker exploration. **A)** Schematic of nasal brush swab sampling from the nasal septum. **B)** Immunohistochemical characterization of the dorsoposterior region of the nasal septum including olfactory neural epithelium (OE). **C)** qPCR analysis of neuronal marker expression in nasal brush swab samples, lymphoblasts (LB), and peripheral blood mononuclear cells (PBMCs). *MS4A1* (lymphoblast marker) and *SPP2* (liver marker) served as negative controls. **D)** Immunocytochemical characterization of nasal brush samples. **E)** Representative single-cell Western blotting (scWB) images identifying cells across the olfactory neuronal lineage. **F)** Quantification of olfactory neuronal lineage cells based on scWB analysis. For each panel, error bars represent mean ± SEM. ****p < 0.0001. Keratin 5 (KRT5), horizontal basal cell marker; βIII-tubulin, immature olfactory neuron marker; doublecortin (DCX), immature olfactory neuron marker; olfactory marker protein (OMP), mature olfactory neuron marker; keratin 18 (KRT18), sustentacular cell markers.

Because nasal brush samples are heterogeneous, methods that enable neuron-specific molecular profiling are required. To address this challenge, we developed a platform that integrates nasal brush swabs with scWB, enabling the quantification of proteins and post-translational modifications at single-cell resolution. We first assessed the compatibility of frozen samples by comparing scWB data generated from paired fresh and frozen aliquots of the same nasal brush sample, given that sample viability during biobanking is crucial for translational pipelines. Nerve growth factor receptor (NGFR), a relatively structurally stable protein^15^, was used as a target for this analysis. Fresh and frozen samples exhibited comparable NGFR signal intensity distributions, indicating that frozen nasal brush samples are suitable for scWB analysis (**Fig. S1A**). We then evaluated whether scWB could resolve the cellular composition of nasal brush swab samples. Consistent with the immunostaining results (**Fig. 1B, D**), scWB identified KRT5-positive horizontal basal cells, doublecortin (DCX)-positive immature olfactory neurons, OMP-positive mature olfactory neurons, and KRT18-positive sustentacular cells (**Fig. 1E, F**). Together, these findings establish a robust and translatable platform for neuron-specific protein profiling of heterogeneous nasal brush samples.

We next applied our platform to AD to determine whether it could detect neuronal AD biomarkers from non-invasive nasal brush samples. We first evaluated phosphorylated tau at Threonine 181 (pT181), a well-established AD biomarker^5,16^, in 20 healthy controls (HC), 15 individuals with mild cognitive impairment (MCI), and 32 patients with clinically diagnosed AD (**Fig. 2A**). Age, sex, and race were comparable across the three groups (**Fig. S2**). HEK293 cells transiently transfected with Tau and CDK5 were used as a positive control for pT181 detection via our scWB platform (**Fig. S3**). pT181 was quantified specifically in OMP-positive mature olfactory neurons (**Fig. 2B, C**). Neuronal pT181 signal intensity was significantly increased in patients with AD compared with both HC and MCI (**Fig. 2C, D**; **Fig. S4**) and was significantly associated with cognitive impairment measured by the Montreal Cognitive Assessment (MoCA) (**Fig. 2E**). Furthermore, we observed a significant difference between the HC and MCI groups (**Fig. 2D**). We next examined phosphorylated tau at threonine 217 (pT217), another established biomarker for AD^5,17^. However, despite testing two independent antibodies, pT217 could not be reliably detected in mature olfactory neurons under the current experimental conditions (**Fig. 2F, G**).

**Figure 2.**
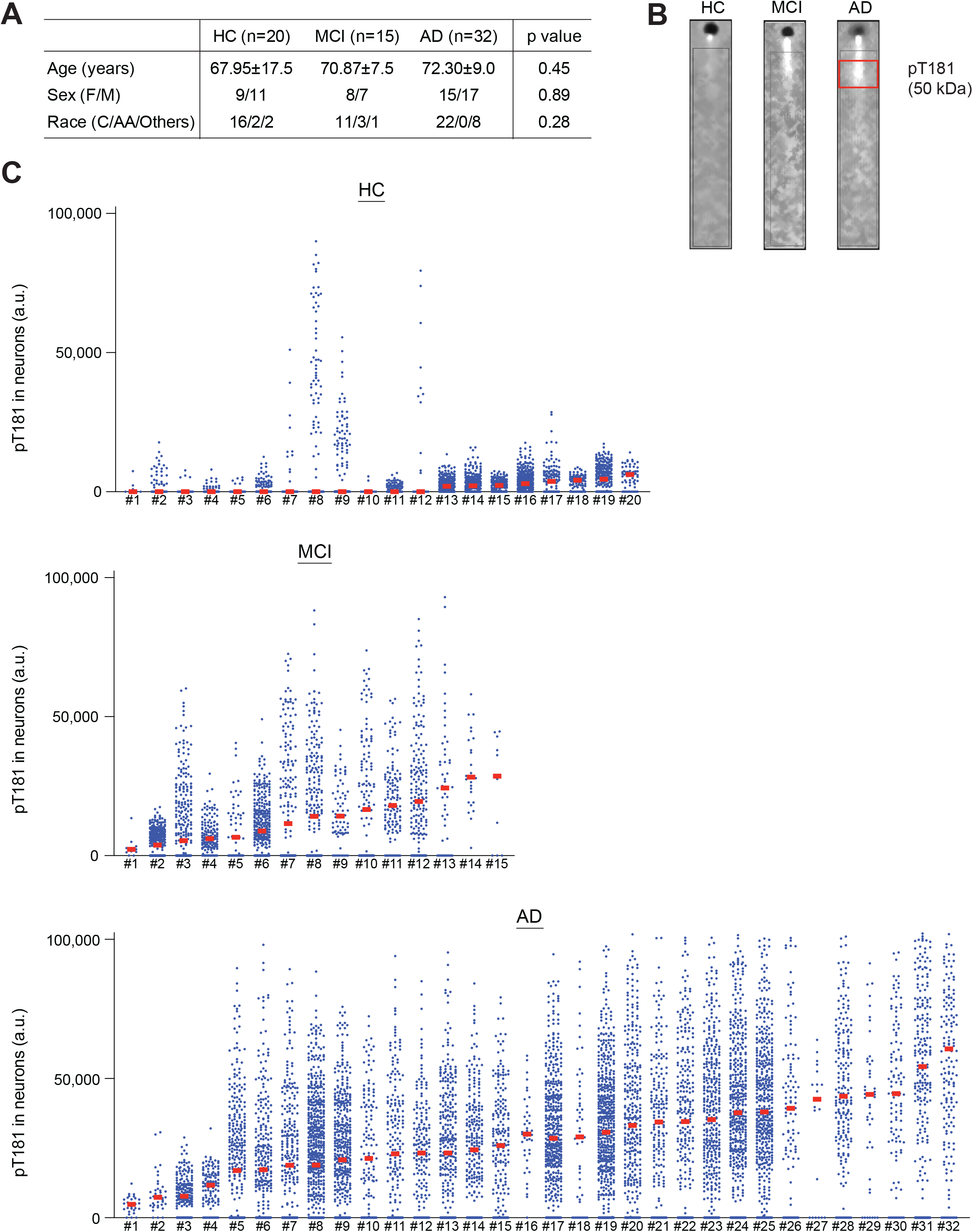
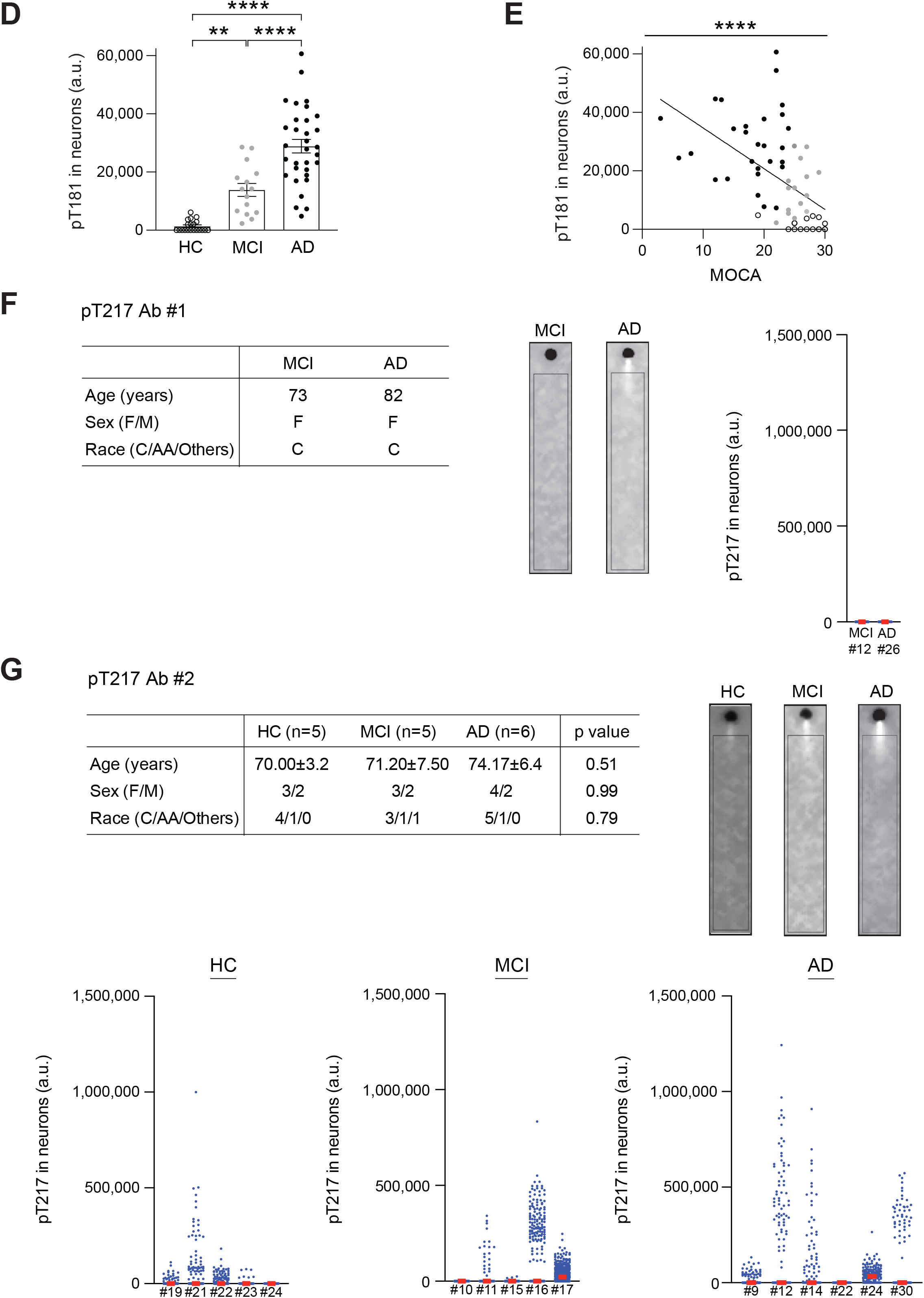
Neuronal phosphorylated tau protein at threonine 181 (pT181) is elevated and associated with cognitive decline in patients with Alzheimer’s disease (AD) and subjects with mild cognitive impairment (MCI). **A)** Demographic characteristics of participants included in the neuronal pT181 analysis. **B)** Representative scWB images of pT181. **C)** Single-cell distribution of neuronal pT181 for each participant, with each data point representing an individual neuron. **D)** Quantification of neuronal pT181 levels across healthy controls (HC), subjects with MCI, and patients with AD. **E)** Correlation between neuronal pT181 levels and cognitive performance measured by the Montreal Cognitive Assessment (MoCA). **F)** Representative scWB images, single-cell distribution, and quantification of neuronal pT217 using Antibody #1. **G)** Representative scWB images, single-cell distribution, and quantification of neuronal pT217 using Antibody #2. For each panel, error bars represent mean ± SEM. **p < 0.01, ****p < 0.0001.

To compare neuronal and plasma pTau within the same individuals, plasma pTau levels were measured using the SIMOA system. In plasma, both pT181 and pT217 levels were significantly elevated in AD compared to HC (**Fig. 3A**) and were associated with cognitive decline (**Fig. 3B**). Plasma pT181 and pT217 levels were also positively correlated with each other (**Fig. 3C**). Notably, pT217 showed a larger difference between AD and HC and a stronger association with cognitive decline than pT181 (**Fig. 3A, B)**. Crucially, however, while the nasal neuronal assay successfully distinguished the MCI group from HC (**Fig. 2D**), the plasma assays failed to do so (**Fig. 3A**). To further validate the potential superiority of the neuronal assays over the plasma assays, we next compared the performance of these two measurements. Neuronal pT181 levels were correlated with both blood pT181 and pT217 (**Fig. 3D**). Importantly, receiver operating characteristic (ROC) analysis showed that neuronal pT181 achieved a higher area under the curve (AUC) than plasma-based measurements, and this difference was significant according to the DeLong test (**Fig. 3E**). The fold-change difference between AD and HC groups was larger for neuronal pT181 than for plasma pTau (**Figs. 2D, 3F**). Altogether, our data with neuronal pT181 measurements highlight the superior diagnostic capacity of the nasal neuronal platform, segregating each group (AD, MCI, and HC) much more clearly.

**Figure 3.**
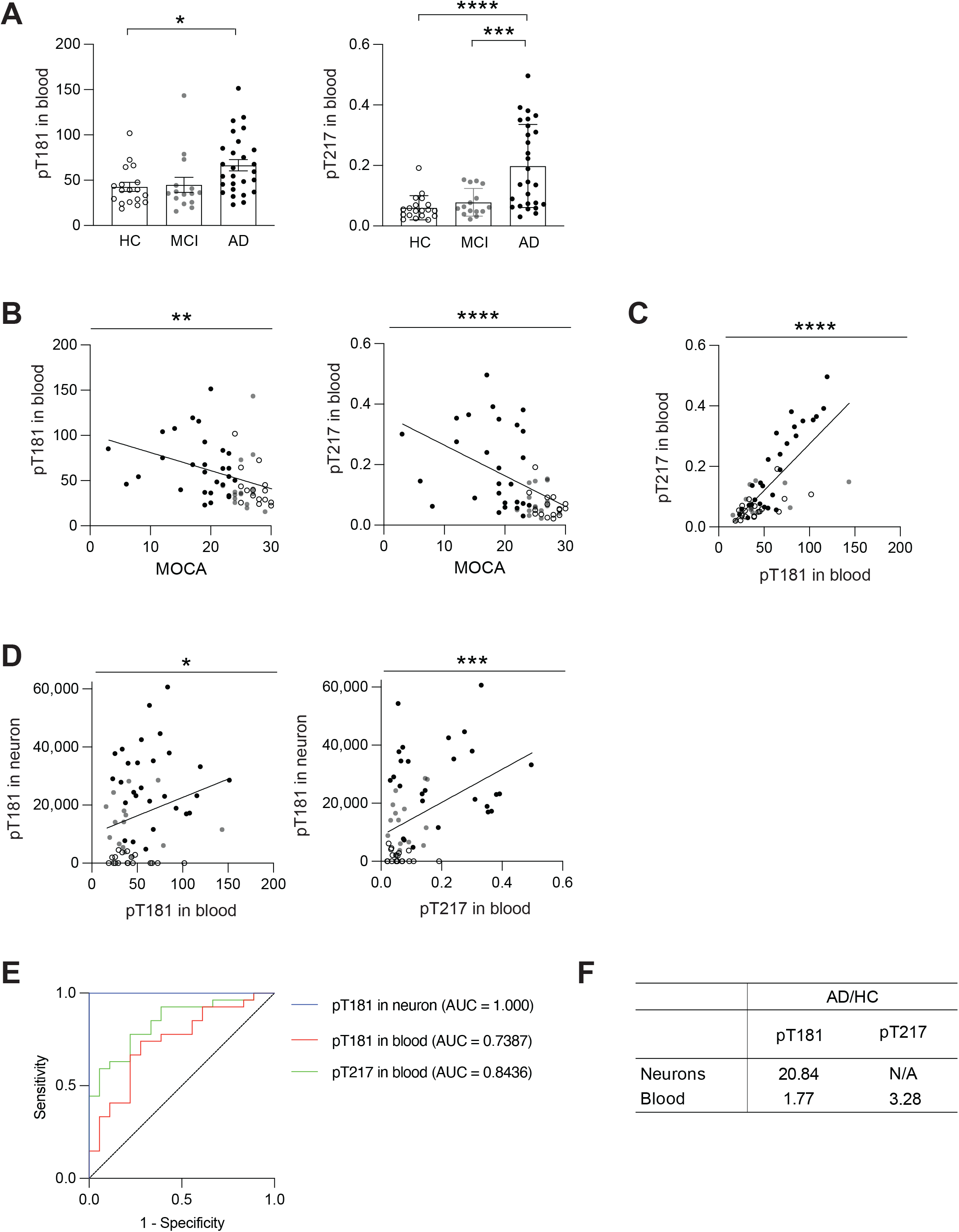
Neuronal pTau shows superior diagnostic performance for AD than plasma pTau. **A)** Quantification of plasma pT181 and pT217 levels in HC, MCI, and AD groups. **B)** Association of plasma pT181 and pT217 levels with cognitive performance measured by MoCA. **C)** Correlation between plasma pT181 and pT217 levels. **D)** Correlation of neuronal pT181 with plasma pT181 and plasma pT217. E) Receiver operating characteristic (ROC) curves comparing the diagnostic performance of neuronal pT181, plasma pT181, and plasma pT217. **F)** Effect size comparison among neuronal pT181, plasma pT181, and plasma pT217. For each panel, error bars represent the mean ± SEM. *p < 0.05, **p < 0.01, ***p < 0.001, ****p < 0.0001.

Finally, we aimed to extend the applicability of our platform and performed a pilot assessment of additional biomarkers for other neurodegenerative disorders. We examined TAR DNA-binding protein 43 (TDP-43)^18^ and phosphorylated α-synuclein^18^ in nasal brush samples. Both biomarkers were detected in a subset of AD patients (**Fig. S5A, B**). Further studies in larger and disease-specific cohorts will be required to fully characterize their disease-associated changes.

In the present study, we developed a non-invasive olfactory platform that enables neuron-specific protein and post-translational modification analyses at single-cell resolution in living patients. As a proof of concept, we applied this platform to test how effectively it can distinguish among patients with AD, MCI subjects, and the HC group, capturing the progressive molecular shifts between these stages. We demonstrated that neuronal pT181 distinguished these three groups, respectively, and exhibited superior diagnostic performance to plasma pTau measurements, highlighting the potential value of directly assessing molecular changes in living neurons. The superiority demonstrated in nasal sampling may be associated with its direct measurement of neuronal proteins, such as pTau, without dilution of the neuronal changes^19^ and free from the contamination of peripheral confounding factors, such as clearance mechanisms^4^.

Nasal brushing has emerged as a valuable and minimally invasive approach for obtaining olfactory neurons from living patients. Recent studies have used single-cell and single-nucleus RNA sequencing to characterize the cellular composition of nasal brush/surgery samples and demonstrate their potential for identifying disease-associated transcriptional changes^2,20,21^, providing important insights into disease biology. Our platform complements these transcriptomic approaches and fundamentally advances them by extending nasal-brush-based analyses to the protein and post-translational modification levels at single-cell resolution. As proteins are the direct mediators of cellular function, whose biological significance is distinctively high, this approach bridges transcriptomic discovery with functionally actionable neuronal targets. Moreover, the minimally invasive nature of nasal brushing makes the platform well suited for repeated sampling and longitudinal monitoring^2^. As disease-modifying therapies for AD are most effective when initiated early^22^, accessible neuron-specific biomarkers obtained through this platform may facilitate early diagnosis, patient stratification, therapeutic monitoring, and evaluation of treatment response.

This study has several limitations. Participant classification was based on clinical diagnosis without confirmation by PET imaging or CSF biomarkers. The cohort size was relatively small, and the cross-sectional design precluded assessment of longitudinal biomarker changes. Future studies in larger, PET- or CSF-confirmed longitudinal cohorts will be important to establish the clinical utility of these neuronal biomarkers. Despite these limitations, our platform provides a unique capability for direct protein and post-translational modification analyses in living human neurons and establishes a foundation for developing functionally actionable neuronal targets across a broad spectrum of brain disorders.

### Study approval

This study was conducted with the approval of the Johns Hopkins Medicine Institutional Review Boards. All study participants provided written informed consent.

## Acknowledgments

This study is supported by National Institutes of Health Grants P50MH136297 (AS) and R21AG070754 (KI), and Subsidies for Current Expenditures to Private Institutions of Higher Education from the Promotion and Mutual Aid Corporation for Private Schools of Japan (KI). The authors wish to extend their gratitude to the participants in the current study. The authors thank Dr. Sandra Lin for conducting the nasal biopsy.

