## Supplementary information for "Olfactory neuroepithelium biomarker platform: a proof from Alzheimer’s disease"

#### Methods

##### Reagents

| Antibodies | Usage | Source | Identifier |
|---|---|---|---|
| KRT5 | scWB, IHC, ICC | BioLegend | Cat#905504 |
| OMP | scWB | aviva | Cat# ARP56669_P050-Biotin |
| OMP | IHC, ICC | FUJIFILM | Cat#544-10001-WAKO |
| Phosphor-Tau (Thr181) | scWB, ICC | invitrogen | Cat#MN1050 |
| KRT18 | scWB, IHC, ICC | invitrogen | Cat#PA5-143130 |
| TDP-43 | scWB | proteintech | Cat#10782-2-AP |
| Phosphor-Apha-synuclein | scWB | abcam | Cat#AB168381 |
| Doublecortin | scWB, IHC, ICC | R&D Systems | Cat#AF10025 |

| Other reagents and equipment | Source | Identifier |
|---|---|---|
| Antigen retrieval buffer | abcam | Cat#ab93648 |
| DMEM/F12 | gibco | Cat#11330-032 |
| FBS | gibco | Cat#A5670701 |
| Penicillin Streptomycin | gibco | Cat#15140-122 |
| Accutase | Sigma-Aldrich | Cat#A6964 |
| ACK lysing buffer | ThermoFisher | Cat# A1049201 |
| Standard scWest Kit | proteinsimple | Cat#K600 |
| Milo | proteinsimple | Cat#P100 |
| InnoScan 710 microarray scanner | INNOPSYS | |
| Cytology brush | Medical Packaging | Cat# CYB-1 |
| 70 mm filter | pluriSelect | Cat#43-50030-03 |

##### Study participants

This study was approved by the Johns Hopkins University School of Medicine Institutional Review Board and all study participants provided written informed consent. Clinically diagnosed patients with Alzheimer's disease (AD), subjects with mild cognitive impairment (MCI), and cognitively normal healthy controls (HC) were recruited at the Johns Hopkins Schizophrenia Center (JHSZC) in collaboration with the Johns Hopkins Alzheimer's Disease Research Center (JHADRC) and MedStar Harbor Hospital. Cognitive functions of all the participants were evaluated by the Montreal Cognitive Assessment (MoCA).

##### Nasal brush swab and single-cell preparation

Nasal brush swabs were performed under endoscopic control at the Johns Hopkins Sinus Center clinic using a cytology brush in awake subjects. Topical nasal oxymetazoline/tetracaine spray was used as a local anesthetic. If sinonasal disease such as mass, edema, nasal polyps, or infection is observed, such subjects are excluded from the study. Based on best visualization of the olfactory cleft in the dorsoposterior region of the nasal septum, a cytology brush was gently positioned in the olfactory cleft and rotated briefly to collect surface cells. The brush was then submerged into collection solution (DMEM/F12 supplemented with 10% FBS and 1%

antibiotics) and transferred to the lab for the single-cell preparation. Nasal brush swab samples were washed in PBS, spun, resuspended in Accutase, and incubated for 10 min at room temperature (RT). Then samples were dissociated mechanically by pipetting into single-cell solution, and 9 times volume of collection solution was added into the cell solution. The cell solution was then filtered through a 70 μm filter and centrifuged for 3 min at 200 × g. The cells were washed in PBS, spun, resuspended in ACK buffer, and incubated for 5 min at RT. After the incubation, the same volume of PBS with 2% FBS was added into the cell solution, and the cell solution was centrifuged for 5 min at 500 × g. The cells were resuspended in collection solution, and the cell number is counted. Finally, the cells were stored in freezing solution (90% FBS and 10% DMSO) at -80℃ until their analytical usage.

**Single-cell western blotting (scWB)**

scWB was performed using the standard scWestern Kit (ProteinSimple) according to the manufacturer's instructions. Briefly, scWest chips were hydrated in Suspension Buffer for at least 10 min. Frozen samples were thawed and resuspended at a concentration of $1.5 \times 10^5$ cells/mL in Suspension Buffer and loaded onto each chip. After 15 minutes to allow cell settling, chips were gently washed three times with Suspension Buffer and examined under a light microscope to assess cell loading efficiency. Optimal loading (10–15% well occupancy) ensures that fewer than 2% of wells contained more than one cell. Cell-loaded chips were subjected to lysis (10 s), electrophoresis (240 V for 60 s), and UV capture (240 s) using the Milo (ProteinSimple). Chips were then incubated with primary antibodies for 2 h at RT, followed by incubation with fluorescent secondary antibodies for 1 h at RT. Chips were washed, dried, and scanned using an InnoScan 710 microarray scanner (Innopsys). For stripping and reprobing, chips were incubated in stripping buffer at 60°C for 90 min, followed by four washes (15 minutes each) in Washing Buffer. Chips were then re-probed with primary antibodies and processed as described above. Images from each spectral channel were analyzed using Scout software (ProteinSimple).

**Immunohistochemistry**

Olfactory epithelium (OE) tissues were obtained by nasal biopsy at the Johns Hopkins Sinus Center clinic, as reported previously[7]. The tissue was immediately fixed in 4% PFA for 8 h at 4 °C, equilibrated sequentially in 20% and 30% sucrose, then embedded in Optimum Cutting Temperature for sectioning. Antigen retrieval was performed in HistoVT at 70°C for 30 min and cooled down to RT. After washing with PBS and permeabilized with 0.5% Triton X-100 in PBS for 10 min at RT, sections were blocked with buffer (10% normal donkey serum and 0.5% Triton X-100 in PBS) for 1 h at RT. Sections were incubated with primary antibodies in dilution buffer (5% normal donkey serum and 0.5% Triton X-100 in PBS) for 2 h at RT. Following three washes in PBS, slides were then incubated with secondary antibodies, each diluted 1:400 for 1 h at RT. Sections were air-dried and mounted with mounting medium. Fluorescent images were acquired using a confocal microscope.

**Immunocytochemistry**

Cells from single-cell suspensions of nasal brush swab samples were fixed with 4% paraformaldehyde for 15 min at RT and seeded onto slides. After antigen retrieval (100X Tris-EDTA Buffer, 70°C for 30 minutes), cells were permeabilized with PBS containing 0.1% Triton X-100 for 5 min at RT. Cells were then blocked with blocking solution (PBS containing 0.2% Triton X-100, 2% bovine serum albumin, and 10% normal donkey serum) for 30 min at RT.

Primary antibodies were applied and incubated for 1 h at RT. Following three washes in PBS, cells were incubated with appropriate secondary antibodies for 1 h at RT. Cells were then air-dried and mounted with mounting medium. Fluorescent images were acquired using a confocal microscope.

**Real-time quantitative reverse transcription PCR (qRT-PCR)**
qRT-PCR of single-cell suspensions of nasal brush samples was performed. Total RNA was isolated using the RNeasy Plus Mini Kit (QIAGEN, 74136) and reverse-transcribed with the High-Capacity RNA-to-cDNA Kit (Applied Biosystems, 4387406). qRT-PCR was performed using TaqMan Gene Expression Assays and TaqMan Fast Advanced Master Mix (Applied Biosystems, 4444557) on a QuantStudio 5 Real-Time PCR System (Thermo Fisher Scientific), following established protocols. All reactions were run in duplicate. The relative mRNA level of target genes was normalized to ACTB and calculated using the $2^{-\Delta\Delta Ct}$ method.

**Transfection**
HEK293 cells were seeded one day before transfection to achieve approximately 70–80% confluency. MYC-tagged Tau and HA-tagged CDK5 plasmids were co-transfected using Lipofectamine 3000 (Thermo Fisher Scientific) according to the manufacturer's protocol. Cells were collected 48 h post-transfection for protein analysis by single-cell western blotting.

**Blood pT181 and pT217 assay**
Plasma pT181 and pT217 were measured with Single Molecule Array (SIMOA) system at Johns Hopkins Bayview Medical Center.

**Statistical analysis**
Statistical analyses were performed using R software (version 4.5.3) and Prism. Group comparisons for continuous variables were conducted using unpaired t-tests or one-way analysis of variance (ANOVA) with Sidak's multiple comparisons test, as appropriate. Linear regression models were used to assess group differences and associations between variables, adjusting for age, sex, and race. Pearson correlation analysis was used to evaluate the associations between continuous variables. Receiver operating characteristic (ROC) curves were generated to assess diagnostic performance, and the area under the curve (AUC) was calculated. Probability values (p values) $< 0.05$ were considered to be statistically significant (*$p < 0.05$, **$p < 0.01$, ***$p < 0.001$, ****$p < 0.0001$).

### Extended Data Fig. 1

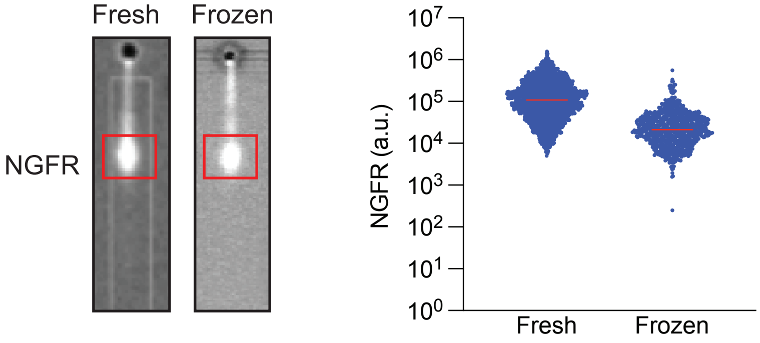


**Extended Data Figure 1.** Sample compatibility of single-cell Western Blotting (scWB). Representative scWB images and quantification of nerve growth factor receptor (NGFR) levels in paired fresh and frozen nasal brush samples from the same participant.

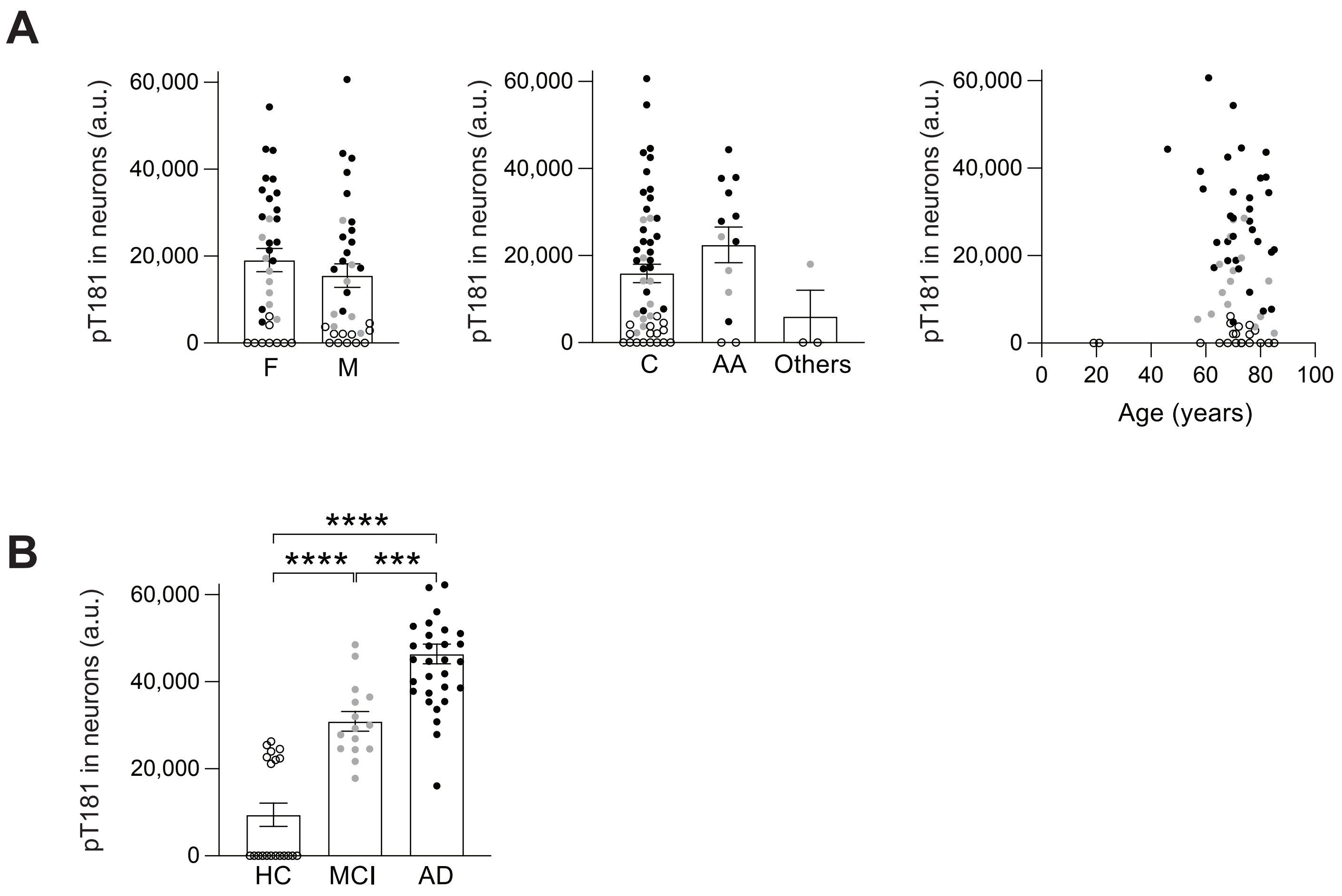


**Extended Data Figure 2.** Neuronal pT181 analysis after the adjustment for confounding factors.
**A)** Neuronal pT181 levels were not associated with sex, race, and age. **B)** Neuronal pT181 analysis data was not changed after the adjustment for confounding factors (sex, race, and age) from the data before the adjustment (see **Fig. 2D**).

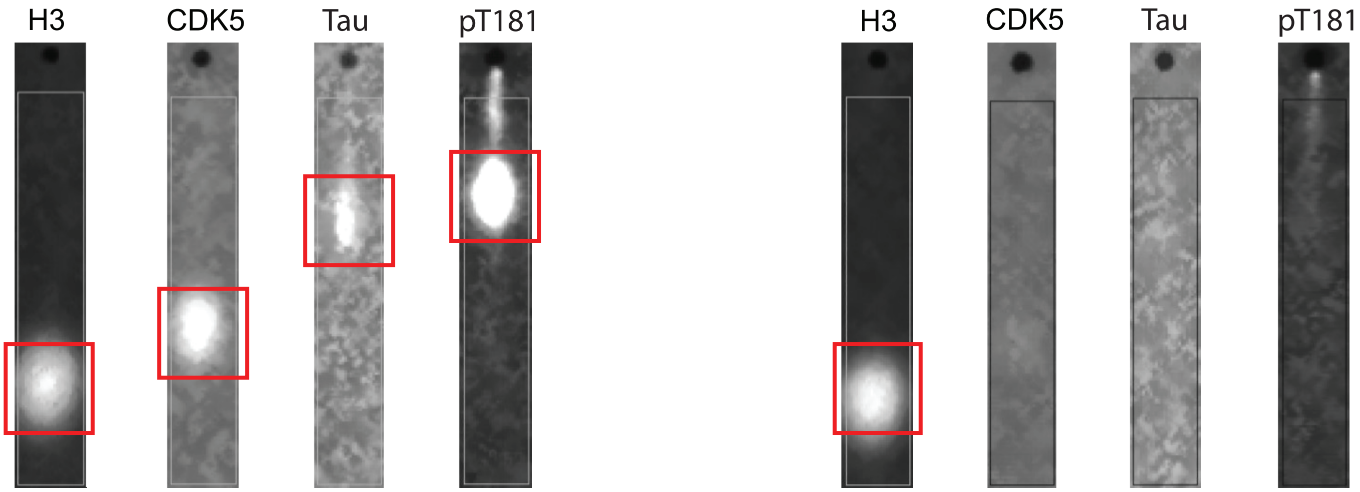


**Extended Data Figure 3.** Positive control for neuronal pT181 detection by scWB. Representative scWB images of pT181 in HEK293 cells co-transfected with human Tau and cyclin-dependent kinase 5 (CDK5) and in non-transfected control cells.

### Extended data Fig. 4

**A**

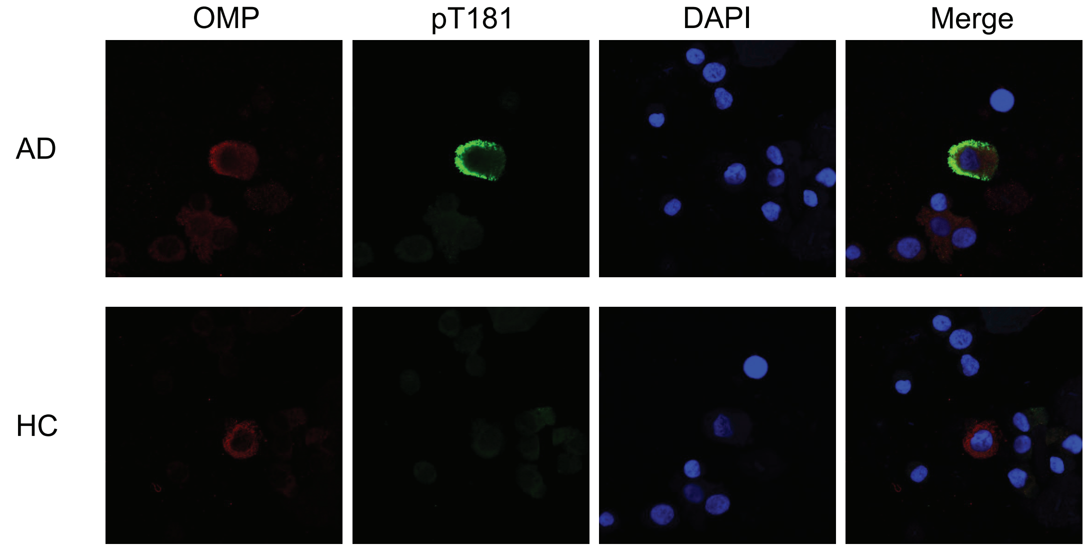


**B**

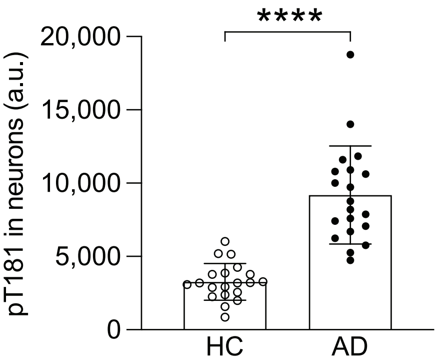


**Extended Data Figure 4.** Immunocytochemical analysis of pT181 in nasal brush samples.
**A)** Representative immunocytochemistry images of pT181 signals in nasal brush samples. **B)** Quantification of pT181 immunofluorescence intensity in HC and patients with AD. Olfactory marker protein (OMP), mature olfactory neuron marker.

### Extended data Fig. 5

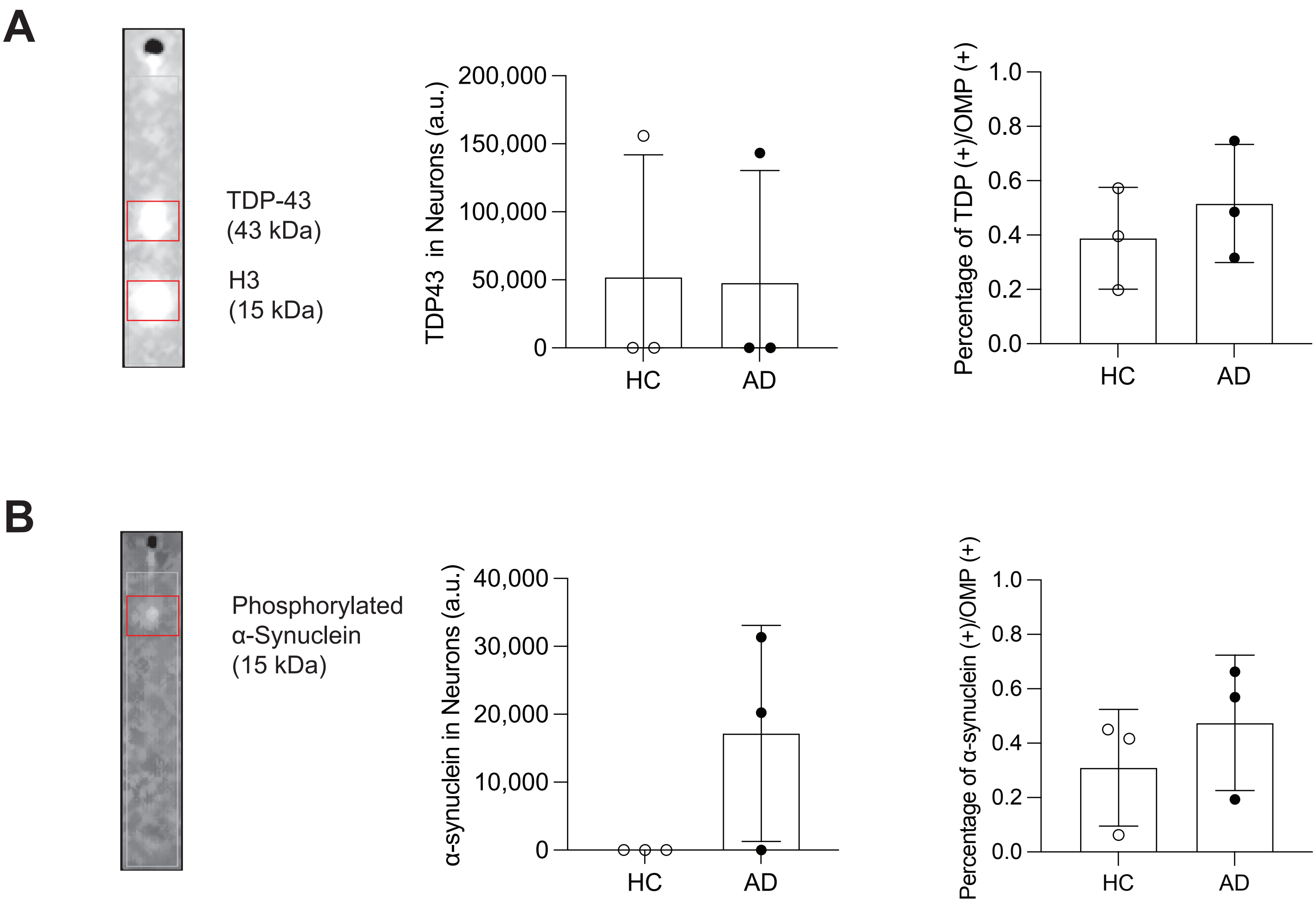


**Extended Data Figure 5.** Detection of additional neurodegenerative biomarkers using the nasal brush swab conbined with scWB. **A)** Representative scWB images and quantification of neuronal TAR DNA-binding protein 43 (TDP-43) levels in HC and patients with AD. **B)** Representative scWB images and quantification of neuronal phosphorylated α-synuclein levels in HC and patients with AD.
